# A multi-omic analysis of optineurin proteinopathy in a yeast model suggests the involvement of lipid metabolism in Amyotrophic Lateral Sclerosis

**DOI:** 10.1101/605998

**Authors:** Daniel M. Bean, Silvia Hnatova, Michael Mülleder, Sandra Magalhães, Daniel J. H. Nightingale, Kathryn S. Lilley, Markus Ralser, Alexandra Nunes, Brian J. Goodfellow, Stephen G. Oliver

## Abstract

Amyotrophic Lateral Sclerosis (ALS) is an incurable fatal neurodegenerative disease for which the precise mechanisms of toxicity remain unclear despite some significant advances in our understanding of the underlying genetic basis. A holistic, integrated view of cellular changes will be critical to understanding the processes of neurodegeneration and the development of effective treatments. Mutant forms of optineurin (a ubiquitin-binding protein involved in autophagy, membrane trafficking, and NF-κB activation) are found associated with cytoplasmic inclusions containing TDP43 or SOD1 in some ALS patients. We have taken a multi-omics approach to understand the cellular response to OPTN overexpression in a yeast model of ALS. We found that genetic interaction screens and metabolomics provided parallel, highly complementary data on OPTN toxicity. Genetic enhancers of OPTN toxicity in yeast relate directly to the native function of OPTN in vesicular trafficking and intracellular transport, suggesting the human OPTN protein is functional when expressed in yeast even though there is no yeast ortholog. Crucially, we find that the genetic modifiers and the metabolic response are distinct for different ALS-linked genes expressed in yeast. This lends strong support to the use of yeast as a model system and omics platform to study ALS.

## Introduction

Amyotrophic Lateral Sclerosis (ALS) is a fatal neurodegenerative disease for which there is no effective treatment available. ALS has been divided into familial (FALS) and sporadic (SALS) forms on the basis of family history, with FALS patients accounting for 5% of all ALS cases overall, but this dichotomy is now questioned[1]. Despite our incomplete picture of the genetic landscape of ALS, it is considered a genetic disease. Associated genetic variants, particularly the hexanucleotide repeat expansion of C9orf72[2,3] and mutations in SOD1[4], TDP43[5,6] and FUS[7,8], are the basis for experimental models of ALS in most model systems. Genetic variants and model organism studies implicate a wide range of cellular pathways in the neurodegenerative processes occurring in ALS, including oxidative stress, RNA metabolism, protein aggregation and degradation (autophagy, and the ubiquitin-proteasome system), and intracellular trafficking[9]. The hallmark histopathological feature of ALS is the presence of intracellular protein aggregates. In most cases, these aggregates contain the TDP43 protein, even though mutations in the TDP43 gene are only a rare cause of ALS. A notable exception is patients with SOD1 mutations, where intracellular aggregates contain the SOD1 protein, but not TDP43. With such a complex pathology underpinning ALS, it is vital to develop multi-omics approaches to understand how the interaction of multiple pathways is driving disease progression.

The OPTN gene encodes Optineurin, a ubiquitin-binding protein involved in autophagy[10–12] membrane trafficking[13] and NF-κB activation[14]. ALS-linked OPTN mutations were first detected in a Japanese cohort of ALS patients[15], where OPTN protein was also found colocalized in cytoplasmic inclusions containing TDP43 or SOD1, suggesting OPTN is broadly involved in ALS regardless of the underlying mutation. Subsequent studies indicated that that OPTN mutations are relatively more common in Asian populations[15–19], and more rare in Caucasian ALS patients[20–23]. Mutations in OPTN are also linked to primary open-angle glaucoma[24] and Paget’s disease of bone[25–27], suggesting that OPTN itself is a key driver of toxicity. Compared to other ALS-linked proteins, there is relatively little research focused on OPTN-ALS. We therefore chose to focus our study on OPTN specifically. The intracellular pathways in which OPTN is involved are strongly conserved between yeast and human cells[28,29]; however, there is no yeast ortholog of optineurin.

Two previous studies have used a yeast model to study OPTN[30,31]. In the first such study, Kryndushkin et al. found that both wild-type and mutant OPTN formed intracellular aggregates and were toxic to yeast[31]. More recently, Jo et al. performed a screen for yeast single-gene deletions that modify the toxicity of human OPTN in yeast[30]. Their screen used a high-copy plasmid to express OPTN, resulting in a level of toxicity that is too high to reliably detect enhancer phenotypes (as the authors also note). Our study builds on this previous work in two main ways. First, we expressed the OPTN coding sequence from a low copy-number CEN plasmid, to get lower toxicity levels and enable the detection of both suppressors and enhancers. Secondly, we also carried out a metabolomic screen to build up a more detailed multi-omic picture of OPTN-ALS.

## Results

### Expression of wild-type human OPTN is toxic in yeast

In line with similar studies[30–35], we cloned human OPTN into yeast expression vectors under the control of the GAL promoter to allow rapidly-inducible strong expression of the transgene. We first confirmed previous results which showed OPTN expression is toxic and that the protein forms aggregates when expressed in yeast[30,31]. Spot tests and liquid culture growth assays both demonstrated the reduced growth of cells expressing OPTN-YFP vs YFP controls (Figure 1). Microscopy demonstrated that OPTN-YFP formed intracellular aggregates in yeast, whereas YFP alone remained diffuse in the cytoplasm (Figure 1). As found previously[31], OPTN tended to aggregate as a single point, whereas TDP-43 and FUS formed multiple diffuse aggregates (not shown).

**Figure 1.**
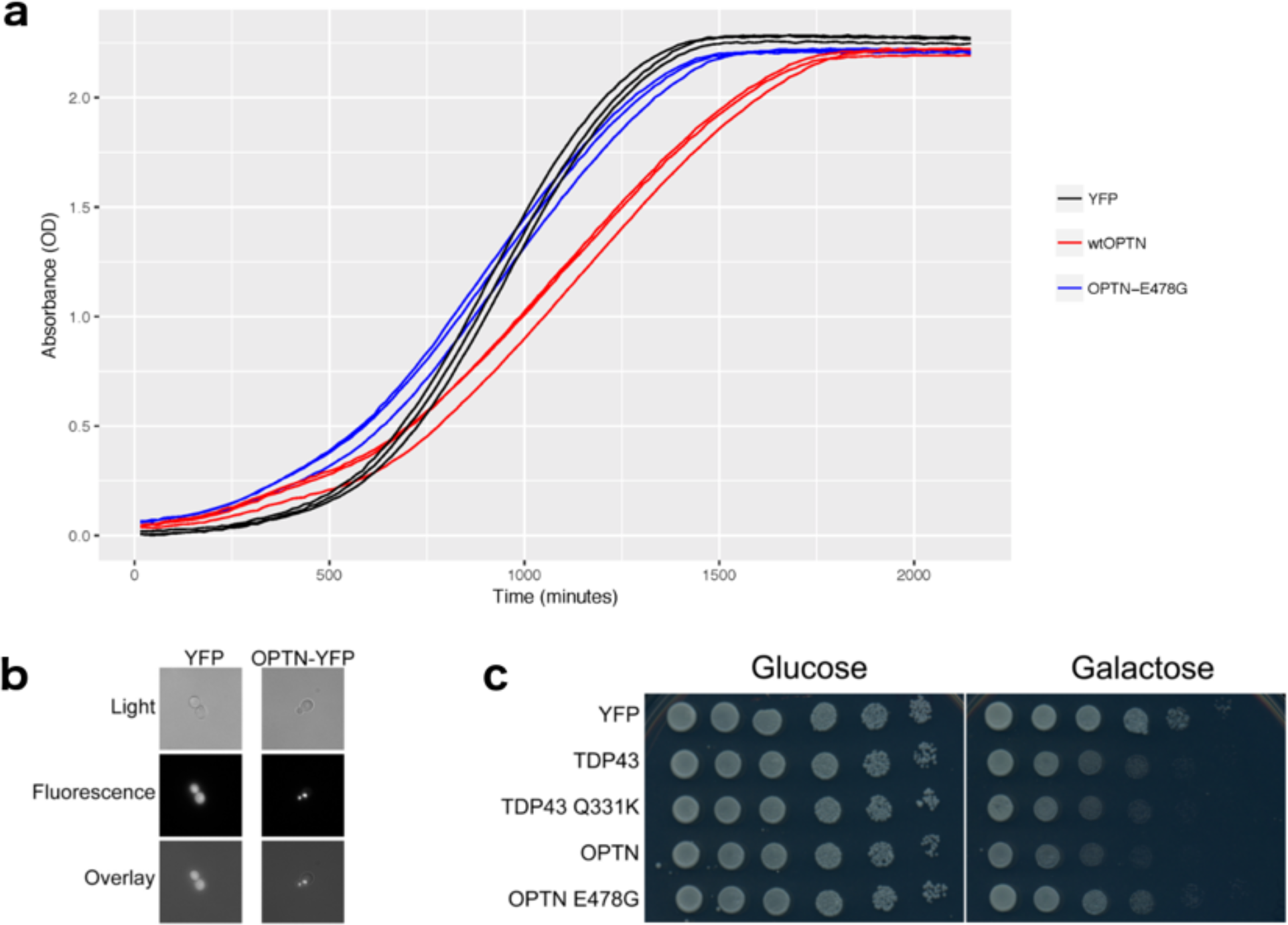
Expression of OPTN is toxic to yeast. a) Growth in liquid culture showed a reduced maximum exponential growth rate for cells expressing OPTN (red) compared to YFP (black). There was no phenotype for cells expressing OPTN-E478G (blue). b) Fluorescence microscopy showed diffuse cytoplasmic fluorescence for YFP alone, whereas OPTN-YFP formed focal aggregates. There was no observable fluorescence for OPTN-E478G with the same exposure (not shown). c) Spot-tests comparing growth on Glucose (expression off) to Galactose (expression on) showed a growth phenotype for OPTN that was similar to TDP43, and a weaker phenotype for OPTN-E478G.

Although there appeared to be a growth phenotype for the E478G mutant in the spot test, this was not confirmed in the liquid culture. Furthermore, the fluorescence of the YFP tag was not detectable for OPTN-E478G-YFP under the same microscopy conditions or in a Western blot. We therefore used only the wild-type OPTN construct for all experiments in this study.

### Genetic screening identifies protective genes

Genetic modifiers of OPTN toxicity were identified using high-throughput synthetic genetic array (SGA) screens[36] to introduce the OPTN-YFP or control plasmids into the BY4741 deletion library. Screens were carried out in biological triplicate, each of which also contained 4 technical repeats. Hits from the screen were defined as strains whose interaction score was greater than 2 standard deviations away from the mean interaction score in all 3 biological repeats, with at least 3 of 4 technical repeats scored and normal growth on SGlu.

We identified 30 suppressors and 64 enhancers of the OPTN growth phenotype (Supplementary Table S1). The only enriched GO term for the suppressors of OPTN toxicity was cytoplasmic translation, suggesting that these genes are not specific suppressors of OPTN but instead affect the expression of OPTN from the plasmid. We therefore focused on the enhancers of OPTN toxicity. Deletion of these enhancer genes increases OPTN toxicity, which suggests their protein products exert a protective effect when present.

The enriched GO terms for the enhancers are shown in Table 1. The most enriched process was “mitochondrion-ER membrane tethering”, which is known to be a key regulator of cell death processes[37]. Several of the enriched terms relate to intracellular trafficking, specifically ER-to-Golgi, which is striking as OPTN plays a major role in Golgi transport and membrane trafficking processes in human cells. Of the 64 suppressors, 52 have at least one human ortholog. The human orthologs are also enriched for vesicle-mediated transport pathways, including ER-to-Golgi transport and Golgi vesicle transport, which suggests the screen results could be translatable to human cells.

**Table 1.**
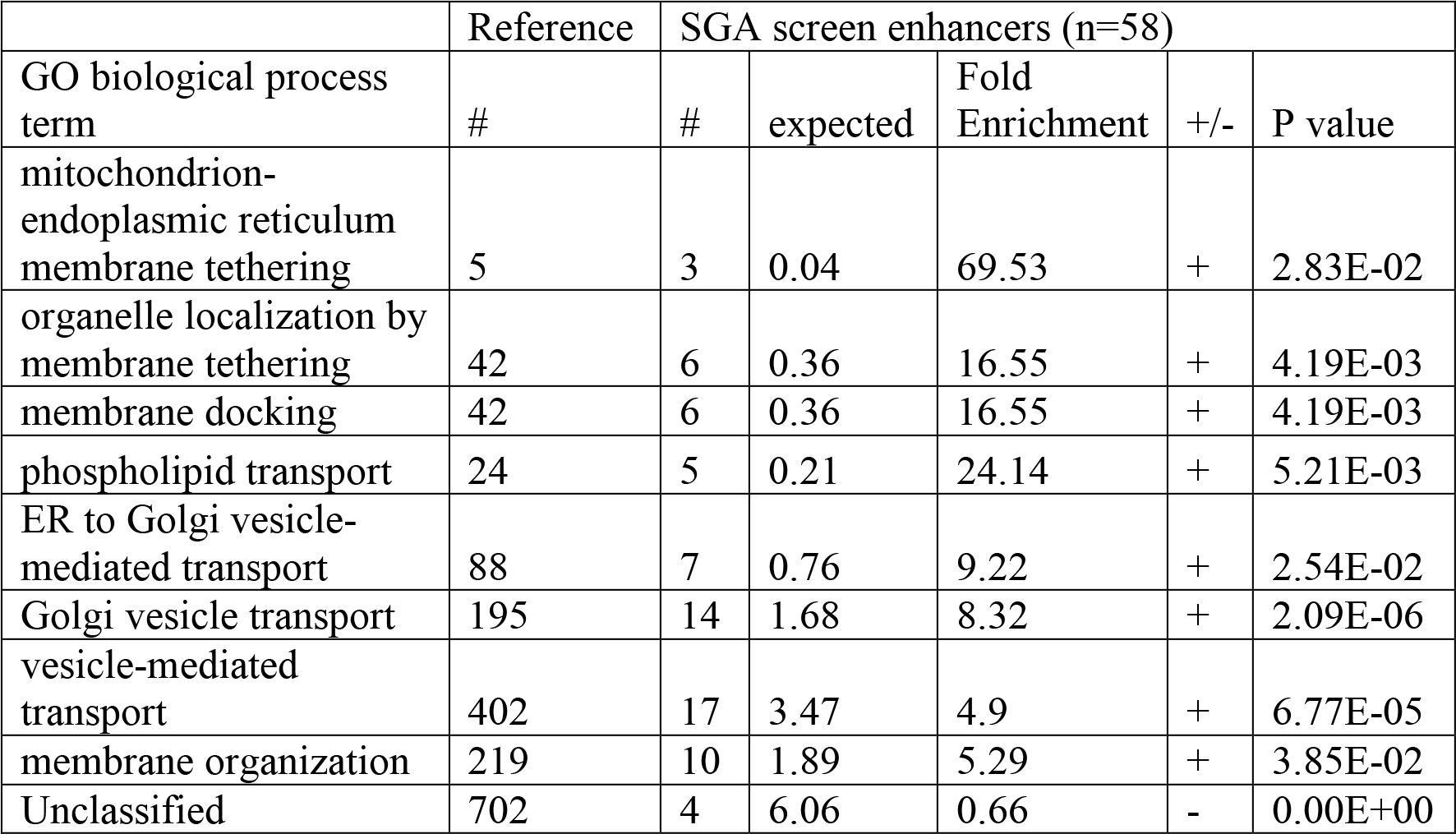
GO biological process term enrichment for enhancer hits. Enrichment of terms for the 58/64 enhancers that were mapped in the PANTHER database. The +/− column indicates enrichment (+) or depletion (−) of the corresponding term in the enhancers gene set. Enrichment calculated with PANTHER release 20170413 and GO release 2017-10-24. P values are shown after applying Bonferroni correction for multiple testing.

|  | Reference | SGA screen enhancers (n=58) |  |  |  |  |
| --- | --- | --- | --- | --- | --- | --- |
| GO biological process term | # | # | expected | Fold Enrichment | +/- | P value |
| mitochondrion-endoplasmic reticulum membrane tethering | 5 | 3 | 0.04 | 69.53 | + | 2.83E-02 |
| organelle localization by membrane tethering | 42 | 6 | 0.36 | 16.55 | + | 4.19E-03 |
| membrane docking | 42 | 6 | 0.36 | 16.55 | + | 4.19E-03 |
| phospholipid transport | 24 | 5 | 0.21 | 24.14 | + | 5.21E-03 |
| ER to Golgi vesicle-mediated transport | 88 | 7 | 0.76 | 9.22 | + | 2.54E-02 |
| Golgi vesicle transport | 195 | 14 | 1.68 | 8.32 | + | 2.09E-06 |
| vesicle-mediated transport | 402 | 17 | 3.47 | 4.9 | + | 6.77E-05 |
| membrane organization | 219 | 10 | 1.89 | 5.29 | + | 3.85E-02 |
| Unclassified | 702 | 4 | 6.06 | 0.66 | - | 0.00E+00 |

Jo et al. recently performed a related OPTN screen in which human OPTN was expressed from a high copy plasmid and genetic interactions were measured using pooled barcode sequencing[30]. Their detected 127 suppressors of toxicity they identified, none is significant as a suppressor or enhancer in our screen (enhancers were unreliable). However, several of the same pathways were implicated by the hits in both studies, particularly lipid metabolism and vesicle-mediated transport.

### Network analysis of protective genes

Given the coherent set of enriched GO terms for these genes, we expected that they, or their products, were likely to genetically or physically interact. Indeed 52/64 of the protective genes interact genetically, and 41/64 of their protein products interact physically (Figure S1). The genetic interaction network is a single component with 232 edges, whereas there are 4 separate components in the protein-protein interaction (PPI) network (containing 32, 4, 3 and 2 proteins, respectively). Together, the genetic and protein interactions connect 55/64 protective genes as a single component with 279 edges. The Syntaxin-like t-SNARE, TLG2 has high centrality (degree, betweeness, and closeness) in both the genetic and protein interaction networks. TLG2 is the yeast ortholog of mammalian Syntaxin-16, which functions in the ER-Golgi vesicle-mediated transport pathway. It is possible that deletion of TLG2 enhances the toxicity of OPTN in yeast by impairing trafficking to the vacuole, thus limiting protein degradation via this pathway. However, previous work in HEK cells found the ubiquitin-proteasome system to be the primary pathway responsible for degradation of OPTN[38].

### Genetic modifiers of OPTN toxicity have little overlap with TDP43 or FUS modifiers in yeast

To determine whether the hits from our screen are specific to OPTN or simply represent a generic response to the expression of a toxic transgene, we compared our results to those of previous SGA screens for TDP43[35] (8 suppressors, 6 enhancers)and FUS[33] (36 suppressors, 24 enhancers) in yeast. These genetic modifiers have limited overlap, consistent with the relevance of our hits to the endogenous function of OPTN. All genes identified as modifiers in more than one screen are shown in Table 2.

**Table 2.** Genetic modifiers identified for multiple ALS genes in yeast. OPTN modifiers identified in this paper, TDP43 modifiers from Armakola et al.[35], FUS modifiers from Sun et al.[33].

| Effect | ALS genes | Deletion systematic name | Deletion standard name | Human ortholog |
| --- | --- | --- | --- | --- |
| Enhancer | OPTN, FUS | YHR030C | SLT2 | MAPK7 |
| Enhancer | OPTN, FUS | YDR148C | KGD2 | DLST |
| Enhancer | OPTN, FUS | YNL052W | COX5A | COX4I1, COX4I2 |
| Enhancer | TDP43, FUS | YML009C | MRPL39 | MRPL33 |
| Suppressor | OPTN, FUS | YBL027W | RPL19B | RPL19 |
| Suppressor | OPTN, FUS | YDR382W | RPP2B | RPLP2 |

The only overlapping genetic modifiers for OPTN were with FUS, with 3 common enhancers and 2 common suppressors. Deletion of the MAP kinase gene SLT2 enhanced the toxic phenotype of both OPTN and FUS, potentially due to dysregulation of peroxisome assembly or the unfolded protein response, both of which are regulated by Slt2p. In a recent study, Jo et al. found that pharmacological inhibition of MAP2K5, an upstream regulator of MAPK7, is a potential target for ALS therapy[30]. The 2 other common enhancers, KGD2 and COX5A, encode mitochondrial proteins involved in the TCA cycle (KGD2) and the inner mitochondrial membrane electron transport chain (COX5A). Both deletions cause a severe reduction in growth rate and may be false positives as the additional effect of OPTN is small.

Two suppressor deletions (rpl19b and rpp2b) were common to the OPTN and FUS screens. Both genes encode ribosomal proteins and therefore affect cytoplasmic translation. It is therefore possible that the protective phenotype is due to lowered expression of the toxic transgene rather than a specific interaction. Finally, despite their more similar cellular functions, only MRPL39, was identified (as an enhancer) in both the TDP43 and FUS screens.

### MS and NMR identification of altered metabolites and lipids

Given that the genetic modifiers of OPTN toxicity were distinct from those previously identified for the ALS risk genes FUS and TDP43, we wondered whether strains expressing these human proteins are also metabolically distinct. The stationary phase (72hr) endometabolomes of yeast strains overexpressing OPTN, FUS and TDP43 were therefore compared to controls using untargeted metabolic profiling by NMR and targeted profiling by MS. A PCA analysis of OD- and TSP-normalized and pareto-scaled NMR data matrices indicated that the profiles are, in fact, distinct (Supplementary Figure S2A) and thus the metabolic differences are due to the specific overexpressed protein. The targeted MS analysis of both the aqueous and lipid fractions (Supplementary Figure S2B) from the same samples confirmed this.

The statistically significant metabolites (from MS and NMR data) that were altered in OPTN samples compared to controls were identified and the effect size calculated (Figure 2).

**Figure 2.**
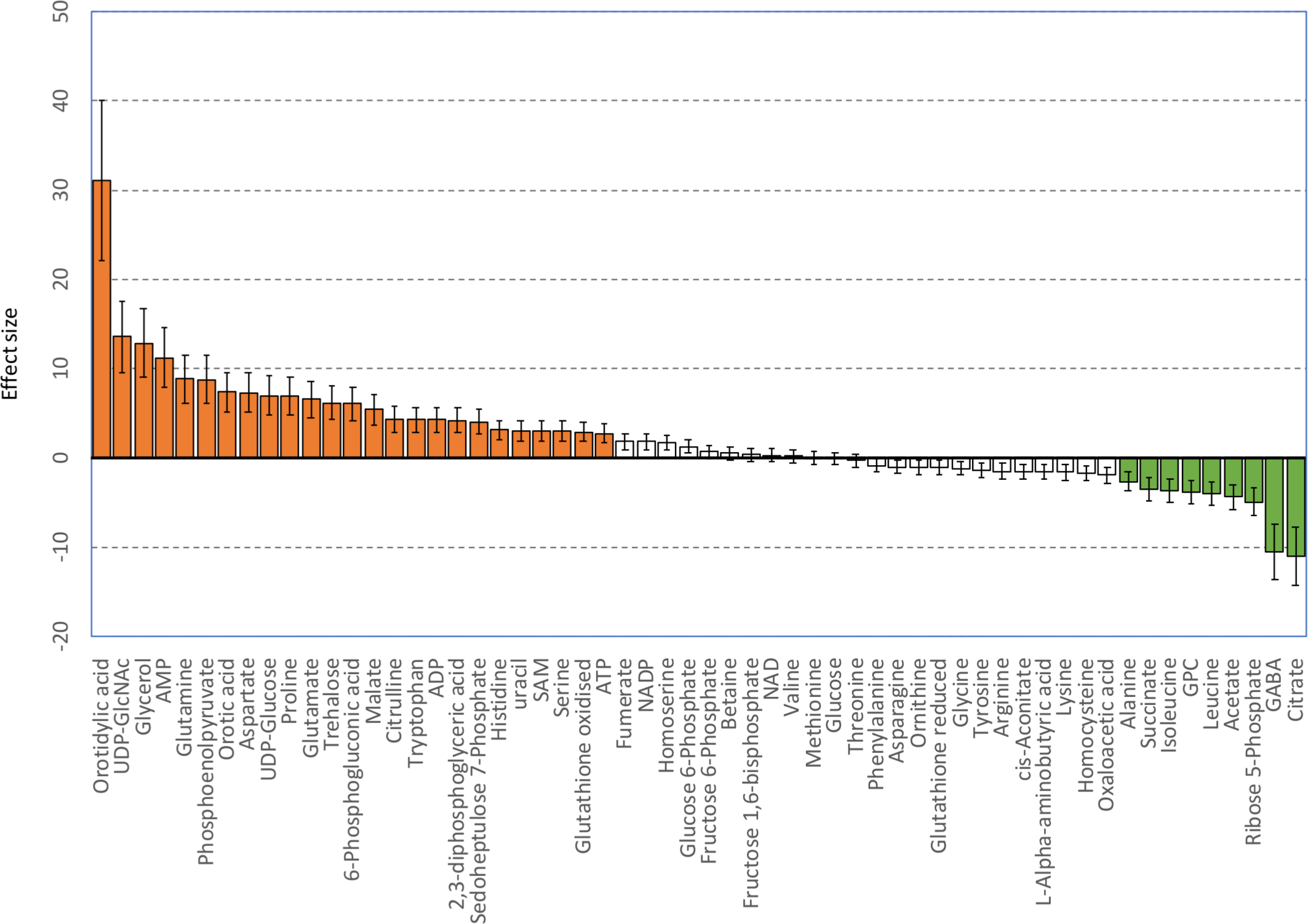
Metabolic ES variations observed for yeast overexpressing OPTN versus controls using data from targeted MS data and untargeted NMR profiles. Statistically significant differences (p<0.05) are shown as orange and green bars.

Metabolites whose levels increase when OPTN is overexpressed include those associated with cellular and metabolic stress - such as γ-aminobutyric acid (GABA), oxidized glutathione, glycerol, and trehalose. Many others are related to cellular energy processes such as the TCA cycle, glycolysis and gluconeogenesis (phosphoenolpyruvate, glutamine; Gln, glutamic acid, adenine nucleotides, and malate). The most significantly increased metabolite was orotidylic acid followed by uridine-diphosphate-N-acetylglucosamine (UDP-GlcNAc) and glycerol, while the metabolites that decreased include the basic amino acids leucine and isoleucine; the TCA cycle intermediates, succinate and citrate; and glycerophosphocholine (GPC).

Metaboanalyst 4.0[39] was used to identify the pathways most affected by the presence of overexpressed OPTN compared to controls (Figure 3). The pathways affected that have the highest impact include: alanine, aspartate and glutamate metabolism; glycine, serine and threonine metabolism; arginine and proline metabolism and glutathione metabolism. Pathways with high significance but lower impact include: butanoate metabolism; pyrimidine/purine metabolism and glycerolipid metabolism.

**Figure 3.**
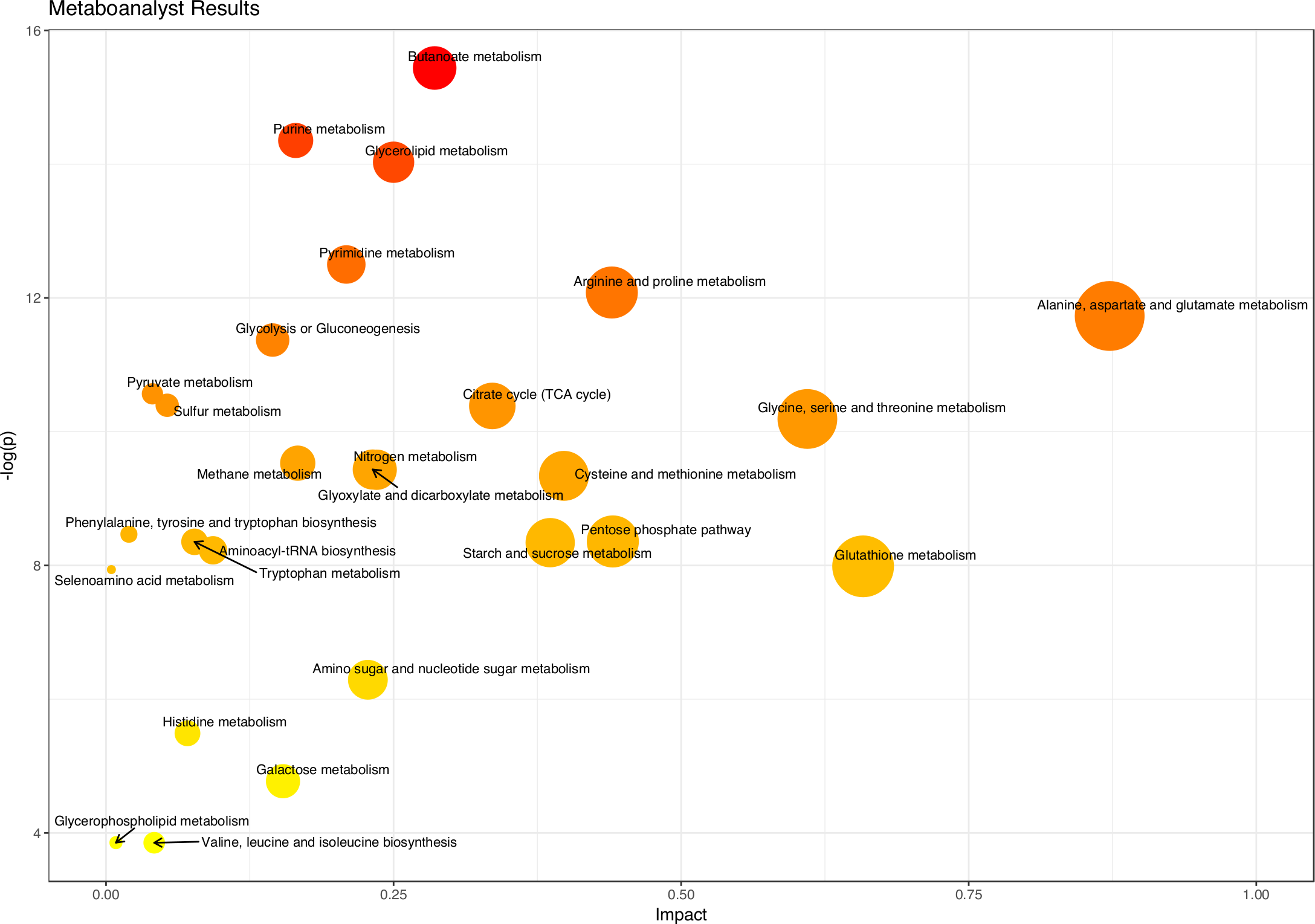
Enriched metabolic pathways observed for yeast overexpressing OPTN versus controls using data from targeted MS data and untargeted NMR profiles in Metaboanalyst 4.0.

To confirm the effect of OPTN on lipid metabolism indicated in the Metaboanalyst results, we also profiled the endometabolome using LC-MS on the organic phase extract. Distinct profiles were again observed (Figure S2B) for OPTN, FUS and TDP43 and 75 (37 negative ion + 38 positive ion) statistically significant m/z species were identified as increased in cells overexpressing OPTN, while 78 (28 negative ion + 50 positive ion) species decreased. Of these, a reduced group was selected based on an effect size above or below 4 and with VIPs larger than 1.56 from a UV-scaled PLS-DA model using the positive ion mode data and ES above or below 4 and VIPS above 1.68 for the negative ion mode data. Identified lipid families are shown in Table 3 and the associated m/z ions and tentative identification in Supplementary Table S2.

**Table 3.** Identified lipid families[63] with differential levels in the LC-MS profiled endometabolome of OPTN yeast vs control.

| Lipid family |  | Increased | Decreased |
| --- | --- | --- | --- |
| FA01 | Fatty acids and conjugates | 3 | 2 |
| ST01 | Sterols | 3 |  |
| GL02 | Diradylglycerols (DAG) | 5 |  |
| GL03 | Triradylglycerols (TAG) | 2 |  |
| GP01 | Glycerophosphocholines | 1 |  |
| GP10 | Glycerophosphates | 1 |  |
| SP02 | Ceramides | 2 |  |
| SP05 | Neutral glycosphingolipids | 1 |  |
| Unknown | - | 2 |  |

## Discussion

In this study, we have generated genome-wide genetic interaction data and carried out a metabolomic screen on the same OPTN yeast model. Significantly, we show that in both profiles, yeast cells expressing OPTN are distinct from yeast cells expressing other human genes associated with ALS (TDP-43 and FUS). We have also identified genetic modifiers of OPTN toxicity that are directly related to the endogenous function of OPTN in human cells.

The wild-type OPTN protein was toxic when expressed in yeast, consistent with related studies of other ALS genes[30,31,33–35,40]. The ALS-linked OPTN mutant E478G was not efficiently expressed in our system, and therefore we used the wild-type protein for all screens. Although we are therefore unable to model the effect of the specific mutations linked to ALS, a significant part of ALS pathology appears to be a proteinopathy, which is recapitulated in this system. For example, Armakola et al. used wild-type TDP-43 for a genome-wide yeast screen and identified dbr1 as a modifier gene that was validated in human neuronal cell lines[35]. Therefore, whilst this system models only part of the pathology of ALS, previous data suggest that results in this type of system are translatable to human cells.

Crucially, the growth phenotype of this yeast proteinopathy model is not a generic response to an overexpressed transgene, as might have been expected. This is shown by the distinct modifiers identified for the ALS-linked genes FUS, TDP-43 and OPTN in genetic interaction screens and in our metabolic profiling. One possible mechanism for these distinct modifiers is that the expressed proteins retain at least part of their endogenous function when expressed in yeast. Yeast is widely used for recombinant protein production, both for research and commercial applications. Whilst the correct folding of any particular recombinant protein is not guaranteed, and a substrate (and/or cofactor) required to carry out that protein’s molecular function may not be available in yeast, it is possible that the OPTN protein retains its native molecular function when expressed in yeast.

In a seminal work, Kachroo et al.[41] found that almost half of the deletion of 414 essential yeast genes could be complemented (“humanised”) by the expression of their human ortholog. Although yeast does not possess an OPTN ortholog, the function of this human protein may be retained through interaction with conserved components of the same pathways. For example, yeast also does not have any orthologs of the Bcl-2 apoptosis regulator proteins, yet expression of mammalian Bax protein induces cell death[42,43], apparently though a mechanism conserved in human cells[44]. Taken together with the direct relevance of the modifiers we identify in our genetic screen to the endogenous function of OPTN (specifically intracellular trafficking and ER-Golgi transport), and the differing metabolic profiles of FUS, TDP-43 and OPTN, we suggest that the phenotype of yeast expressing transgenic OPTN is due to the native properties or functions of the OPTN protein, and not simply due to its overexpression.

The metabolic profiling results suggest that OPTN is affecting the TCA cycle, which may be related to general mitochondrial dysfunction. As overexpression of a non-native protein in yeast would also be expected to induce the UPR (and ER stress), the increased levels of some amino acids may be a result of this process causing increased protein catabolism, a reduction in protein biosynthesis or a reduction in amino acid utilization/biosynthesis. Oxidative stress was increased in OPTN yeast compared to controls as the GSH_red_/GSH_ox_ ratio is 1.4 times lower in these cells (Supplementary Figure S3). Interestingly, an increase in the amino acid proline is seen in OPTN expressing yeast. As the presence of this amino acid has been found to minimize protein aggregation and the depletion of proline has been linked to the inhibition of the UPR[45], our results are consistent with OPTN aggregation and triggering of the UPR in our yeast system. Increased UDP-GlcNAc levels suggests that cell wall biosynthesis may be decreased, and lipid metabolism also appears to be altered as indicated by a decrease in GPC and an increase in Ser and in UDP-glucose, both involved in sphingolipid biosynthesis. As UDP-GlcNAc is also intimately involved in the production of N-glycans, with biosynthesis first taking place in the ER and subsequently in the Golgi apparatus, any disruption in trafficking between these organelles could also affect UDP-GlcNAc levels. This may corroborate our conclusion that the phenotype of yeast cell expressing OPTN reflects the native function of the protein (ER-Golgi transport) and is not a generic response to an exogenous protein. Our profiling data also indicates that orotidylic acid had the largest increase of all the aqueous assigned metabolites detected in our study. At this time, we do not have an explanation as to the significance of this perturbation.

A number of metabolomic biomarkers have been proposed for neurological diseases including ALS[46–49]. The results from Wuolikainen et al.[46] show some correlation with our results where 3 of the top 5 positively correlated ALS metabolites (Pro, Trp, AMP) in plasma are also seen as increased in our OPTN cells. A deficit in RNA synthesis was also seen, suggesting a decrease in the PPP. The decrease in ribose-5-phosphate seen in our yeast model is consistent with this. Basic amino acids were also indicated as potential biomarkers in CSF and plasma. However, we see decreases in yeast, while increases are seen in human fluids. A recent metabolomic study of a neuronal cellular model of ALS[50] included analysis of metabolite variations seen for cells overexpressing SOD1 and G93A SOD1 under serum deprivation. These results, in general, compare well with those found in our OPTN stationary-phase yeast model.

Our lipid data from LC-MS profiling indicated that many lipid species were significantly increased in OPTN-expressing cells (16 increasing above ES 7 compared to 2 decreasing). In fact, only OPTN (and not FUS or TDP43) showed a larger number of increased lipids compared to decreased lipids. Thus, OPTN appears to affect lipid metabolism even as an exogenous protein in yeast. An LC-MS profiling study of the lipidome for the CSF of ALS patients has identified a number of lipid biomarkers for ALS^46^. Phosphatidylcholines, sphingomyelins, glucosylceramides and sterols were found to be increased, while TAG was decreased in ALS patients. We also see increases in lipids from these families in OPTN-overexpressing yeast cells. Although in a different model, two ALS studies using different SOD1 mutated mice[51,52] also demonstrated that lipids such as sphingolipids, ceramides and glucosylceramides are increased in spinal cord fluid and skeletal muscle. Therefore, it appears that OPTN may be producing a yeast phenotype that reflects lipidome effects seen in in vivo situations.

We have studied genetic interactions, integrating those results with protein interaction data, and metabolites. However, additional ‘omics approaches could, and should, be added to build both a broad and deep intracellular understanding of ALS. This panoramic view of ALS is necessary to predict the impact of perturbations to this system either by mutation or, eventually, by treatment.

## Methods

### Yeast strains and media

The Synthetic Genetic Array (SGA)[36] starter strain Y7092 (MATα can1Δ::STE2pr-Sp_his5 lyp1Δ his3Δ1 leu2Δ0 ura3Δ0 met15Δ0) was used for all experiments was mated with the BY4741 deletion library in the SGA. Strains were manipulated, and media prepared using standard microbiological techniques. Yeast were cultured in synthetic minimal media without uracil, and with either 2% glucose (SGlu), 2% raffinose (SRaf) or 2% galactose (SGal) as the carbon source.

To induce expression from the plasmid in liquid culture, single colonies were picked into 50μl SGlu in a 96-well plate and incubated at 30°C with shaking. After 24h, 10μl of the SGlu cultures was inoculated into 200μl SRaf and incubated at 30°C with shaking. After 24h, 20μl of the SRaf cultures were inoculated into 100μl fresh SRaf and incubated at 30°C with shaking. SGal medium was inoculated with a 1:20 dilution of this SRaf culture in either 96-well or 384-well plates.

For induction in the SGA, deletion mutants containing the plasmid were pinned onto SGal plates at 384 colonies per plate. After incubation at 30°C for 24h, these colonies were pinned onto SRaf at 384 colonies per plate and incubated at 30°C for 24h. Each colony from the SRaf plates was pinned onto SGal 4 times (1536 colonies per plate), and incubated at 30°C. Plates were scanned at 300dpi after 48h of growth. All pinning steps were performed using a Singer ROTOR HDA.

### Plasmid construction

OPTN in pEGFP-C3 was kindly donated by Dr. Justin Yerbury. The OPTN ORF was PCR amplified from pEGFP-C3 and cloned into the Gateway donor plasmid pDONR221 (KanR) via the Gateway BP reaction according to the manufacturer’s instructions. The E478G mutation was introduced by site-directed mutagenesis using the Aglient QuikChange II Site-Directed Mutagenesis Kit. Donor plasmids containing FUS and TDP43 were kindly donated by the Gitler lab.

All pDONR221 Gateway donors were cloned into the Gateway destination vector pAG416GAL-ccdb-EYFP[53] (CEN, URA3, AmpR, referred to as pAG416) obtained from Addgene via the LR reaction, according to manufacturer instructions.

The empty pAG416 vector was modified to create the control (YFP only) plasmid by removing the sequence between the GAL1 promoter and YFP. In the unmodified pAG416, YFP is approximately 1750 bp from the GAL1 promoter, resulting in weak expression. The YFP coding sequence was PCR-amplified from pAG416 and, in parallel, pAG416 was digested with Kpn1 and Not1. The larger restriction fragment was gel-purified and recombined with the PCR product in yeast. The resulting plasmid, pAG416-short, was used as the control in all experiments.

All plasmid sequences were confirmed by restriction mapping and DNA sequencing. YFP expression (either alone or as a tag) was confirmed by fluorescence microscopy and Western blot using anti-GFP (AbCam antibody ab6556) and anti-histone H3 (AbCam ab1791) as a loading control. The presence of the expressed protein (OPTN, TDP43 or FUS) was confirmed by in-gel digestion, followed by LC-MS/MS. Briefly, yeast protein extraction for both western blotting and LC-MS/MS was performed according to[54]. Total extract, corresponding to approximately 1.7 × 10^6^ cells, was resolved by SDS-PAGE gel electrophoresis. Following Coomassie staining, a band was excised from the gel that corresponded to a ca. 20 kDa range around the predicted molecular weight of the eYFP-tagged human protein. The band was dissected into cubes of approximately 1 mm and destained (using ammonium bicarbonate), reduced (with dithiothreitol) and alkylated (using iodoacetamide). The sample was digested for 16 h at 37C using a 1:50 (w/w) ratio of Sequencing Grade Modified Trypsin (Promega):protein in the gel. The digest was analysed by LC-MS/MS on a nanoAcquity UPLC system (Waters) coupled in-line to a LTQ Orbitrap Velos mass spectrometer (ThermoFisher Scientific), essentially according to[55], but with the modification that MS2 scans were performed on the twenty most intense ions per survey scan with a charge of 2+ or above. In all cases, the human protein corresponding to the expressed transgene was correctly identified in the LC-MS/MS data.

### Mass spectrometry data processing

Raw mass spectrometry data files were converted to MGF format using MSConvert (version 3.0.9283, Proteowizard). MGF files were searched using an in-house Mascot server (version 2.6.0, Matrix Science) against three databases at the same time, which were a canonical S. cerevisiae database, downloaded from UniProt (March 2017; 6,749 sequences), a canonical isoformal version of the human SwissProt database (November 2016; 42,144 sequences) and the cRAPome database of common mass spectrometry contaminants[56] (January 2017; 115 sequences). Precursor tolerance was set to 20 ppm and fragment tolerance to 0.6 Da. Carbamidomethylation of cysteine was specified as a fixed modification and oxidation of methionine as a variable modification.

### Genetic interaction screen

Genetic interactions were screened using Synthetic Genetic Array technology as described[36], the only modification being the use of URA3 as the selectable marker for the query strain (Y7092 transformed with OPTN-YFP or control plasmid) instead of NatMX4. Interactions were scored from images scanned at 300dpi after 48h of growth on SGal using Gitter[57] in SGAtools[58] (available at http://sgatools.ccbr.utoronto.ca/). Colonies that failed to grow on SGlu media were excluded from the analysis. Hits from the screen were defined as strains greater than 2 standard deviations away from the mean interaction score in all 3 biological repeats with at least 3 of 4 technical repeats scored and normal growth on SGlu.

### Interaction networks

Yeast interaction data was retrieved from the 06/11/2017 update of YeastMine[59] (https://yeastmine.yeastgenome.org) and analysed in Cytoscape[60].

### Quenching and extraction of the endometabolome

For all samples (TDP, FUS, OPTN) and controls two biological replicates were prepared and two technical replicates used. After the final incubation in 25 ml galactose medium (OD600 = 0.05) the samples were harvested at 72 hr and immediately placed on ice. A protocol similar to Palomino-Schätzlein et al. (2013) was followed[61]. Pelleting and resuspension in cold phosphate buffer was carried out followed by centrifugation and flash freezing in liquid nitrogen. For metabolite extraction, 500 μL of methanol/chloroform (2:1) at 4°C was added to the frozen samples and the pellet was resuspended by vortexing after 5 min. Five freeze/thaw cycles of 1 min in liquid nitrogen and 2 min on ice were then carried out followed by addition of 250 μL of chloroform and 250 μL of MilliQ water. Vortexing for 1 min was followed by a 30 min centrifugation (20°C, 16000 × g). The upper aqueous layer was collected carefully with a Gilson pipette. The lipophilic phase was collected into a glass vial. The samples were dried using a stream of N_2_ gas (NitroFlowLab) on a Techne Dri-Block(R) DB30 (aqueous 2-4 hr, lipid 10 min). The aqueous samples were further dried in a speedvac (1-2 hr, Savant Speed Vac(R) SPD111V). All extracts were stored at 4°C.

### NMR spectroscopy

For the NMR analysis, the aqueous extracts were re-suspended in 620 μl D_2_O, 0.01% TSP and 100 mM pH 7.0 phosphate buffer (Na_2_HPO_4_ 100 mM, pH 7.0). All samples were centrifuged (1 min, 20 °C, 16,000 g) before transferring to 5mm NMR tubes. The ^1^H-NMR spectra were recorded at 298K on a Bruker Avance III 500 MHz spectrometer using a TXI or TCI probe. 1D ^1^H spectra were acquired using a NOESY pulse sequence to suppress the water resonance, with a sweep width of 7002 Hz (14 ppm), 32k data points, a recycle delay of 12 s, a mixing time of 100 ms and 128 scans per free induction decay (FID).

### Mass spectrometry

#### Aqueous fractions

Metabolite concentrations for the aqueous fractions were determined on a liquid chromatography (Agilent 1290 Infinity) and tandem mass spectrometry (Agilent 6460) system. All compounds were identified by comparing retention time and fragmentation pattern with analytical standards. The instrument was operated in single reaction monitoring mode. Ion transitions and analytical methods used for metabolite identification and concentration determination are given in Supplementary Tables S3 and S4. Metabolite concentrations were determined by external calibration. Solvents were of UPLC grade and chemicals of at least analytical grade. Specific conditions for amino acids, other polar metabolites and UDP-N-acetylglucosamine are given in supplementary methods.

#### Lipid fractions

Samples, re-suspended in 200μl of HPLC grade methanol, were analysed in both positive and negative ion modes using a Waters Xevo G2 quadrupole time of flight (Q-ToF) combined with an Acquity Ultra Performance Liquid Chromatogram (UPLC) (Waters Corporation, Manchester, UK). Injection volumes and conditions along with gradient parameters and data acquisition are given in supplementary methods.

### Multivariate analysis

iNMR software (http://www.inmr.net) was used to process the NMR spectra with zero filling to 64k data points and 0.3 Hz line broadening being applied before Fourier transformation. The spectra were manually phased, baseline corrected, referenced to TSP at 0.00ppm and exported (0.5-9.5 ppm for aqueous phase, 0.5-8 ppm for lipid phase) as a matrix. The spectra were normalized to OD600 at the time of extraction and then to TSP. The OD600 values at the time of extraction were (mean ± standard deviation): control = 7.98 ± 0.14, TDP43 = 6.80 ± 0.09, FUS = 7.91 ± 0.08 and OPTN = 7.87 ± 0.15. The normalised spectra were then checked in iNMR and, if necessary, alignment was carried out using using the ‘speaq’ package in R. The water region was excluded from the alignment.

MVA were performed using the ropls package[62] in R. Initial Principal Components Analysis identified outliers that were excluded from subsequent analyses. The identification of metabolites for NMR was carried out by comparing the spectra with those of standard compounds from the Biological Magnetic Resonance Data Bank, the Yeast Metabolome Database and the Human Metabolome Database. The relative amounts of the NMR metabolites and the effect size were determined by integrating the area under the most well-separated metabolite peak in iNMR and then using in-house R scripts. MS metabolite concentrations were used directly after normalisation to OD600. Pairwise t-tests were carried out using the False Discovery Rate (FDR) to adjust for multiple testing. Effect sizes were calculated and corrected for small sample sizes using the formula:

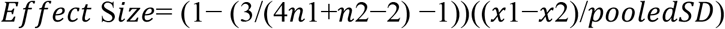

where pooled SD is the pooled standard deviation, x1 and x2 are the mean levels of metabolite x and n1 and n2 are the number of replicates. Metaboanalyst 4.0[39] was used to identify enriched metabolic pathways. The final list of metabolites used included 51 from the targeted MS analysis and 7 (orotidylic acid, glycerophosphocholine, trehalose, glycerol, betaine, uracil and acetate) non-duplicated metabolites from NMR.

For the lipid LCMS data, positive and negative ion mode deisotoped results were normalised to total area and analysed as for aqueous NMR and MS data. Tentative identification of lipids with statistically significant effect size differences was carried out using m/z data and the LIPID MAPS Online Tools[63] (Supplementary Table S2).

## Data availability

The metabolomics data has been deposited at Metabolights with access code, MTBLS796.

## Acknowledgements

The work of D.M.B., G.F., S.H. and S.G.O. was supported by the Wellcome Trust/MRC (grant code: 089703/Z/09/Z). D.J.H.N. was supported by a BBSRC Strategic Longer and Larger grant, awarded to K.S.L. (award BB/L002817/1) The authors are grateful to Dr. Justin Yerbury for the OPTN plasmid, Prof. Aaron Gitler for the TDP-43 and FUS plasmids and Dr. Mike Deery for LC-MS/MS analysis of our plasmid overexpression samples. We thank Steven Murfitt (University of Cambridge) for the acquisition of the LCMS/MS data for the endometabolome lipid fraction samples.

B.J.G. and A.N. were supported by grants UID/BIM/04501/2013 and UID/CTM/50011/2013, co-funded by Fundação para a Ciência e Tecnologia I.P. (PIDDAC) and by European Regional Development Fund (FEDER) and POCI-01- 0145-FEDER-007628 and POCI-01-0145-FEDER-007679, funded by the Operational Programme Competitiveness and Internationalization COMPETE 2020. S.M. is supported by Fundação para a Ciência e Tecnologia I.P. through the individual PhD grant SFRH/BD/131820/2017. The NMR spectrometers used in this work are part of the National NMR Network (PTNMR) and are partially supported by Infrastructure Project N° 022161 (co-financed by FEDER through COMPETE 2020, POCI and PORL and FCT through PIDDAC).

This paper represents independent research part funded by the National Institute for Health Research (NIHR) Biomedical Research Centre at South London and Maudsley NHS Foundation Trust and King’s College London. The views expressed are those of the author(s) and not necessarily those of the NHS, the NIHR or the Department of Health.

## Author contributions

Conceptualisation – DMB, BJG, SGO

Formal Analysis – DMB, BJG

Funding Acquisition – SGO, BJG

Investigation – DMB, MM, SM, SH, DJHN

Supervision – KSL, MR, AN, BJG, SGO

Writing – Original Draft Preparation – DMB, BJG

Writing – Review and Editing – SGO, DMB, BJG, MM, SM, SH, DJHN, KSL, AN

## References

1. Chiò A et al. 2014 Genetic counselling in ALS: facts, uncertainties and clinical suggestions. J. Neurol. Neurosurg. & Psychiatry 85, 478 LP–485.

2. Renton AE et al. 2011 A hexanucleotide repeat expansion in C9ORF72 is the cause of chromosome 9p21-linked ALS-FTD. Neuron (doi:10.1016/j.neuron.2011.09.010)

3. DeJesus-Hernandez M et al. 2011 Expanded GGGGCC Hexanucleotide Repeat in Noncoding Region of C9ORF72 Causes Chromosome 9p-Linked FTD and ALS. Neuron (doi:10.1016/j.neuron.2011.09.011)

4. Halper RDRSTPDFDASPHADDGJORJPDHXRDKAM-YDCAGSMBRTR. et al. 1993 Mutations in Cu/Zn superoxide dismutase gene are associated with familial amyotrophic lateral sclerosis. Nature 362, 59–62. (doi:10.1038/362059a0)

5. Kabashi E et al. 2008 TARDBP mutations in individuals with sporadic and familial amyotrophic lateral sclerosis. Nat. Genet. (doi:10.1038/ng.132)

6. Sreedharan J et al. 2008 TDP-43 mutations in familial and sporadic amyotrophic lateral sclerosis. Science (doi:10.1126/science.1154584)

7. Kwiatkowski TJ et al. 2009 Mutations in the FUS/TLS gene on chromosome 16 cause familial amyotrophic lateral sclerosis. Science (80-.). (doi:10.1126/science.1166066)

8. Vance C et al. 2009 Mutations in FUS, an RNA Processing Protein, Cause Familial Amyotrophic Lateral Sclerosis Type 6. Science (80-.). (doi:10.1126/science.1165942)

9. Hardiman O, Al-Chalabi A, Chio A, Corr EM, Logroscino G, Robberecht W, Shaw PJ, Simmons Z, Van Den Berg LH. 2017 Amyotrophic lateral sclerosis. Nat. Rev. Dis. Prim. 3. (doi:10.1038/nrdp.2017.71)

10. Dikic I et al. 2011 Phosphorylation of the Autophagy Receptor Optineurin Restricts Salmonella Growth. Science (80-.). 333, 228–233. (doi:10.1126/science.1205405)

11. Wong YC, Holzbaur ELF. 2014 Optineurin is an autophagy receptor for damaged mitochondria in parkin-mediated mitophagy that is disrupted by an ALS-linked mutation. Proc. Natl. Acad. Sci. 111, E4439–E4448. (doi:10.1073/pnas.1405752111)

12. Sirohi K, Swarup G. 2016 Defects in autophagy caused by glaucoma-associated mutations in optineurin. Exp. Eye Res. 144, 54–63. (doi:10.1016/j.exer.2015.08.020)

13. Nagabhushana A, Chalasani ML, Jain N, Radha V, Rangaraj N, Balasubramanian D, Swarup G. 2010 Regulation of endocytic trafficking of transferrin receptor by optineurin and its impairment by a glaucoma-associated mutant. BMC Cell Biol. 11. (doi:10.1186/1471-2121-11-4)

14. Zhu G, Wu CJ, Zhao Y, Ashwell JD. 2007 Optineurin Negatively Regulates TNFα- Induced NF-κB Activation by Competing with NEMO for Ubiquitinated RIP. Curr. Biol. 17, 1438–1443. (doi:10.1016/j.cub.2007.07.041)

15. Maruyama H et al. 2010 Mutations of optineurin in amyotrophic lateral sclerosis. Nature 465, 223–226. (doi:10.1038/nature08971)

16. Li C, Ji Y, Tang L, Zhang N, He J, Ye S, Liu X, Fan D. 2015 Optineurin mutations in patients with sporadic amyotrophic lateral sclerosis in China. Amyotroph. Lateral Scler. Front. Degener. 16, 485–489. (doi:10.3109/21678421.2015.1089909)

17. Iida A et al. 2012 Novel deletion mutations of OPTN in amyotrophic lateral sclerosis in Japanese. Neurobiol. Aging 33. (doi:10.1016/j.neurobiolaging.2011.12.037)

18. Iida A, Hosono N, Sano M, Kamei T, Oshima S, Tokuda T, Kubo M, Nakamura Y, Ikegawa S. 2012 Optineurin mutations in Japanese amyotrophic lateral sclerosis. J. Neurol. Neurosurg. Psychiatry. 83, 233–235. (doi:10.1136/jnnp.2010.234963)

19. Naruse H et al. 2012 Mutational analysis of familial and sporadic amyotrophic lateral sclerosis with OPTN mutations in Japanese population. Amyotroph. Lateral Scler. 13, 562–566. (doi:10.3109/17482968.2012.684213)

20. Millecamps S et al. 2011 Screening of OPTN in French familial amyotrophic lateral sclerosis. Neurobiol. Aging 32. (doi:10.1016/j.neurobiolaging.2010.11.005)

21. Johnson L, Miller JW, Gkazi AS, Vance C, Topp SD, Newhouse SJ, Al-Chalabi A, Smith BN, Shaw CE. 2012 Screening for OPTN mutations in a cohort of British amyotrophic lateral sclerosis patients. Neurobiol. Aging 33. (doi:10.1016/j.neurobiolaging.2012.06.023)

22. Del Bo R et al. 2011 Novel optineurin mutations in patients with familial and sporadic amyotrophic lateral sclerosis. J. Neurol. Neurosurg. Psychiatry 82, 1239–1243. (doi:10.1136/jnnp.2011.242313)

23. Belzil V V., Daoud H, Desjarlais A, Bouchard JP, Dupré N, Camu W, Dion PA, Rouleau GA. 2011 Analysis of OPTN as a causative gene for amyotrophic lateral sclerosis. Neurobiol. Aging 32. (doi:10.1016/j.neurobiolaging.2010.10.001)

24. Rezaie T et al. 2002 Adult-Onset Primary Open-Angle Glaucoma Caused by Mutations in Optineurin — Supplemental Data. Science 295, 1077–1079. (doi:10.1126/science.1066901)

25. Albagha OME et al. 2010 Genome-wide association study identifies variants at CSF1, OPTN and TNFRSF11A as genetic risk factors for Paget’s disease of bone. Nat. Genet. 42, 520–524. (doi:10.1038/ng.562)

26. Michou L, Conceição N, Morissette J, Gagnon E, Miltenberger-Miltenyi G, Siris ES, Brown JP, Cancela ML. 2012 Genetic association study of UCMA/GRP and OPTN genes (PDB6 locus) with Paget’s disease of bone. Bone 51, 720–728. (doi:https://doi.org/10.1016/j.bone.2012.06.028)

27. Silva IAL, Conceição N, Gagnon É, Caiado H, Brown JP, Gianfrancesco F, Michou L, Cancela ML. 2018 Effect of genetic variants of OPTN in the pathophysiology of Paget’s disease of bone. Biochim. Biophys. Acta - Mol. Basis Dis. 1864, 143–151. (doi:https://doi.org/10.1016/j.bbadis.2017.10.008)

28. Bonifacino JS, Glick BS. 2018 The Mechanisms of Vesicle Budding and Fusion. Cell 116, 153–166. (doi:10.1016/S0092-8674(03)01079-1)

29. Finley D, Ulrich HD, Sommer T, Kaiser P. 2012 The ubiquitin-proteasome system of Saccharomyces cerevisiae. Genetics 192, 319–360. (doi:10.1534/genetics.112.140467)

30. Jo M et al. 2017 Yeast genetic interaction screen of human genes associated with amyotrophic lateral sclerosis: Identification of MAP2K5 kinase as a potential drug target. Genome Res. 27, 1487–1500. (doi:10.1101/gr.211649.116)

31. Kryndushkin D, Ihrke G, Piermartiri TC, Shewmaker F. 2012 A yeast model of optineurin proteinopathy reveals a unique aggregation pattern associated with cellular toxicity. Mol. Microbiol. 86, 1531–1547. (doi:10.1111/mmi.12075)

32. Couthouis J et al. 2011 A yeast functional screen predicts new candidate ALS disease genes. Proc. Natl. Acad. Sci. U. S. A. 108, 20881–90. (doi:10.1073/pnas.1109434108)

33. Sun Z, Diaz Z, Fang X, Hart MP, Chesi A, Shorter J, Gitler AD. 2011 Molecular determinants and genetic modifiers of aggregation and toxicity for the als disease protein fus/tls. PLoS Biol. 9. (doi:10.1371/journal.pbio.1000614)

34. Ju S et al. 2011 A Yeast Model of FUS/TLS-Dependent Cytotoxicity. PLoS Biol. 9, e1001052. (doi:10.1371/journal.pbio.1001052)

35. Armakola M et al. 2012 Inhibition of RNA lariat debranching enzyme suppresses TDP-43 toxicity in ALS disease models. Nat. Genet. 44, 1302–1309. (doi:10.1038/ng.2434)

36. Baryshnikova A, Costanzo M, Dixon S, Vizeacoumar FJ, Myers CL, Andrews B, Boone C. 2010 Synthetic genetic array (SGA) analysis in Saccharomyces cerevisiae and schizosaccharomyces pombe. Methods Enzymol. 470, 145–179. (doi:10.1016/S0076-6879(10)70007-0)

37. Grimm S. 2012 The ER-mitochondria interface: The social network of cell death. Biochim. Biophys. Acta - Mol. Cell Res. 1823, 327–334. (doi:10.1016/j.bbamcr.2011.11.018)

38. Mao J, Xia Q, Liu C, Ying Z, Wang H, Wang G. 2017 A critical role of Hrd1 in the regulation of optineurin degradation and aggresome formation. Hum. Mol. Genet. 26, 1877–1889. (doi:10.1093/hmg/ddx096)

39. Xia J, Sinelnikov I V., Han B, Wishart DS. 2015 MetaboAnalyst 3.0-making metabolomics more meaningful. Nucleic Acids Res. 43, W251–W257. (doi:10.1093/nar/gkv380)

40. Johnson BS, McCaffery JM, Lindquist S, Gitler AD. 2008 A yeast TDP-43 proteinopathy model: Exploring the molecular determinants of TDP-43 aggregation and cellular toxicity. Proc. Natl. Acad. Sci. 105, 6439–6444. (doi:10.1073/pnas.0802082105)

41. Kachroo AH, 1, Laurent JM, Yellman CM, Meyer AG, 1 2, Claus O. Wilke, 1, 2 3, Marcotte EM. 2015 Systematic humanization of yeast genes reveals conserved functions and genetic modularity. Science (80-.). 348, 921–925. (doi:10.1126/science.aaa0769)

42. William G, Christine S, Bhabatosh C. 1999 Role of mitochondria and C-terminal membrane anchor of Bcl-2 in Bax induced growth arrest and mortality in Saccharomyces cerevisiae. FEBS Lett. 380, 169–175. (doi:10.1016/0014-5793(96)00044-0)

43. Xu Q, Reed JC. 2018 Bax Inhibitor-1, a Mammalian Apoptosis Suppressor Identified by Functional Screening in Yeast. Mol. Cell 1, 337–346. (doi:10.1016/S1097-2765(00)80034-9)

44. Manon S, Chaudhuri B, Guérin M. 1997 Release of cytochrome c and decrease of cytochrome c oxidase in Bax-expressing yeast cells, and prevention of these effects by coexpression of Bcl-xL. FEBS Lett. 415, 29–32. (doi:https://doi.org/10.1016/S0014-5793(97)01087-9)

45. Liang X, Dickman MB, Becker DF. 2014 Proline biosynthesis is required for endoplasmic reticulum stress tolerance in Saccharomyces cerevisiae. J. Biol. Chem. 289, 27794–27806. (doi:10.1074/jbc.M114.562827)

46. Wuolikainen A, Jonsson P, Ahnlund M, Antti H, Marklund SL, Moritz T, Forsgren L, Andersen PM, Trupp M. 2016 Multi-platform mass spectrometry analysis of the CSF and plasma metabolomes of rigorously matched amyotrophic lateral sclerosis, Parkinson’s disease and control subjects. Mol. BioSyst. 12, 1287–1298. (doi:10.1039/C5MB00711A)

47. Kori M, Aydin B, Unal S, Arga KY, Kazan D. 2016 Metabolic Biomarkers and Neurodegeneration: A Pathway Enrichment Analysis of Alzheimer’s Disease, Parkinson’s Disease, and Amyotrophic Lateral Sclerosis. Omi. A J. Integr. Biol. 20, 645–661. (doi:10.1089/omi.2016.0106)

48. Blasco H et al. 2017 Lipidomics Reveals Cerebrospinal-Fluid Signatures of ALS. Sci. Rep. 7. (doi:10.1038/s41598-017-17389-9)

49. H. B, F. P, B. MH, H.GP, P. V, R. AC, P. C. 2016 Metabolomics in amyotrophic lateral sclerosis: how far can it take us? Eur. J. Neurol. 23, 447–454. (doi:10.1111/ene.12956)

50. Valbuena GN, Rizzardini M, Cimini S, Siskos AP, Bendotti C, Cantoni L, Keun HC. 2016 Metabolomic Analysis Reveals Increased Aerobic Glycolysis and Amino Acid Deficit in a Cellular Model of Amyotrophic Lateral Sclerosis. Mol. Neurobiol. 53, 2222–2240. (doi:10.1007/s12035-015-9165-7)

51. Dodge JC et al. 2015 Glycosphingolipids are modulators of disease pathogenesis in amyotrophic lateral sclerosis. Proc. Natl. Acad. Sci. (doi:10.1073/pnas.1508767112)

52. Henriques A et al. 2015 Amyotrophic lateral sclerosis and denervation alter sphingolipids and up-regulate glucosylceramide synthase. Hum. Mol. Genet. (doi:10.1093/hmg/ddv439)

53. Alberti S, Gitler AD, Lindquist S. 2007 A suite of Gateway® cloning vectors for high-throughput genetic analysis in Saccharomyces cerevisiae. Yeast 24, 913–919. (doi:10.1002/yea.1502)

54. von der Haar T. 2007 Optimized Protein Extraction for Quantitative Proteomics of Yeasts. PLoS One 2, e1078. (doi:10.1371/journal.pone.0001078)

55. Chen C, Buhl E, Xu M, Croset V, Rees JS, Lilley KS, Benton R, Hodge JJL, Stanewsky R. 2015 Drosophila Ionotropic Receptor 25a mediates circadian clock resetting by temperature. Nature 527, 516–520. (doi:10.1038/nature16148)

56. Mellacheruvu D et al. 2013 The CRAPome: a contaminant repository for affinity purification-mass spectrometry data. Nat. Methods 10, 730–6. (doi:10.1038/nmeth.2557)

57. Wagih O, Parts L. 2014 gitter: A Robust and Accurate Method for Quantification of Colony Sizes From Plate Images. G3&#58; Genes|Genomes|Genetics 4, 547–552. (doi:10.1534/g3.113.009431)

58. Wagih O et al. 2013 SGAtools: One-stop analysis and visualization of array-based genetic interaction screens. Nucleic Acids Res. 41. (doi:10.1093/nar/gkt400)

59. Balakrishnan R, Park J, Karra K, Hitz BC, Binkley G, Hong EL, Sullivan J, Micklem G, Cherry JM. 2012 YeastMine-An integrated data warehouse for Saccharomyces cerevisiae data as a multipurpose tool-kit. Database 2012. (doi:10.1093/database/bar062)

60. Shannon P, Markiel A, Ozier O, Baliga NS, Wang JT, Ramage D, Amin N, Schwikowski B, Ideker T. 2003 Cytoscape: a software environment for integrated models of biomolecular interaction networks. Genome Res. 13, 2498–504. (doi:10.1101/gr.1239303)

61. Palomino-Schätzlein M, Molina-Navarro MM, Tormos-Pérez M, Rodríguez-Navarro S, Pineda-Lucena A. 2013 Optimised protocols for the metabolic profiling of S. cerevisiae by1H-NMR and HRMAS spectroscopy. Anal. Bioanal. Chem. 405, 8431–8441. (doi:10.1007/s00216-013-7271-9)

62. Thévenot EA, Roux A, Xu Y, Ezan E, Junot C. 2015 Analysis of the Human Adult Urinary Metabolome Variations with Age, Body Mass Index, and Gender by Implementing a Comprehensive Workflow for Univariate and OPLS Statistical Analyses. J. Proteome Res. 14, 3322–3335. (doi:10.1021/acs.jproteome.5b00354)

63. Fahy E, Sud M, Cotter D, Subramaniam S. 2007 LIPID MAPS online tools for lipid research. Nucleic Acids Res. (doi:10.1093/nar/gkm324)

